# What does your cell really do? Model-based assessment of mammalian cells metabolic functionalities using omics data

**DOI:** 10.1101/2020.04.26.057943

**Authors:** Anne Richelle, Benjamin P. Kellman, Alexander T. Wenzel, Austin W.T. Chiang, Tyler Reagan, Jahir M. Gutierrez, Chintan Joshi, Shangzhong Li, Joanne K. Liu, Helen Masson, Jooyong Lee, Zerong Li, Laurent Heirendt, Christophe Trefois, Edwin F. Juarez, Tyler Bath, David Borland, Jill P. Mesirov, Kimberly Robasky, Nathan E. Lewis

**Affiliations:** Novo Nordisk Foundation Center for Biosustainability at the University of California, San Diego, School of Medicine, La Jolla, CA 92093, United States; Department of Pediatrics, University of California, San Diego, School of Medicine, La Jolla, CA 92093, United States; Bioinformatics and Systems Biology program, University of California, San Diego, La Jolla, CA, USA 92093, United States; Department of Medicine, University of California, San Diego, School of Medicine, La Jolla, CA 92093, United States; Moores Cancer Center, University of California, San Diego, La Jolla, CA 92093, United States; Department of Bioengineering, University of California, San Diego, La Jolla, CA 92093, United States; Department of Computer Science and Engineering, University of California, San Diego, La Jolla, CA 92093, United States; Luxembourg Centre for Systems Biomedicine, University of Luxembourg, Esch-sur-Alzette, Luxembourg; Department of Biomedical Informatics, UC San Diego Health, University of California, San Diego, La Jolla, CA 92093, United States; Renaissance Computing Institute, The University of North Carolina at Chapel Hill, Chapel Hill, NC 27517, United States; Department of Genetics, University of North Carolina at Chapel Hill, Chapel Hill, NC 27514, United States; School of Information and Library Science, University of North Carolina at Chapel Hill, Chapel Hill, NC 27599, United States; Carolina Health and Informatics Program, University of North Carolina at Chapel Hill, Chapel Hill, NC 27599, United States

**Author notes:** Correspondence: Nathan E. Lewis.

**Keywords:** metabolic function, omic data, systems biology, functional analysis

## Abstract

Large-scale omics experiments have become standard in biological studies, leading to a deluge of data. However, researchers still face the challenge of connecting changes in the omics data to changes in cell functions, due to the complex interdependencies between genes, proteins and metabolites. Here we present a novel framework that begins to overcome this problem by allowing users to infer how metabolic functions change, based on omics data. To enable this, we curated and standardized lists of metabolic tasks that mammalian cells can accomplish. We then used genome-scale metabolic networks to define gene modules responsible for each specific metabolic task. We further developed a framework to overlay omics data on these modules to predict pathway usage for each metabolic task. The proposed approach allows one to directly predict how changes in omics experiments change cell or tissue function. We further demonstrated how this new approach can be used to leverage the metabolic functions of biological entities from the single cell to their organization in tissues and organs using multiple transcriptomic datasets (human and mouse). Finally, we created a web-based CellFie module that has been integrated into the list of tools available in GenePattern (www.genepattern.org) to enable adoption of the approach.

## Introduction

High-throughput omics technologies allow researchers to comprehensively monitor cells and tissues at the molecular level, and record subtle molecular changes that may contribute to the acquisition of a specific phenotype. However, the complex interdependencies between the gene, protein, and metabolite components limit our capacity to identify the molecular basis of specific phenotypic changes. Therefore, it remains challenging to extract tangible biological meaning from omics data.

Many approaches exist to systematically interpret gene expression changes, ranging from simple enrichment analyses to detailed mechanistic systems biology modeling. Several user-friendly approaches have been developed that allow any researcher to test for enrichment in groups of genes, e.g., pathways, biological processes, or ontology terms^1, 2^. Such approaches are invaluable for identifying groups of genes that are more frequently differentially expressed, but the methods are limited in their capacity to describe how the differential changes impact cellular metabolic functions. To interpret the impact on function, mathematical models of pathways can be used. For example, genome-scale metabolic network reconstructions are knowledgebases of all metabolic pathways in an organism^3–5^. These networks directly link genotype to phenotype, since they mathematically describe the mechanisms by which all cell parts (e.g., membranes, proteins, etc.) are concurrently made. Thus, approaches have emerged to analyze omics data in the context of these models^6, 7^, yielding a wealth of detailed insights into the mechanisms underlying complex biological processes^8^. However, these approaches are not widely used since they are quite complex, requiring months of analysis by experts with years of specialized training.

Here, we propose an alternative approach for the interpretation of omics data (e.g., differentially expressed genes) that captures the simplicity of enrichment analyses, while providing mechanistic insights into how differential expression impacts specific cellular functions, based on pre-computed model simulations. To this end, genome-scale metabolic networks were decomposed into many smaller metabolic tasks^9, 10^. We curated and standardized these tasks, resulting in a collection of hundreds of tasks covering 7 major metabolic activities of a cell (energy generation, nucleotide, carbohydrate, amino acid, lipid, vitamin & cofactor and glycan metabolism). We further developed a framework to directly predict the activity of these metabolic functions from transcriptomic data. To this end, we used genome-scale models of mammalian metabolism to define gene modules responsible for the activation of pathways required for each specific metabolic task. Through this platform, users can overlay their data and comprehensively quantify the propensity of a cell line or tissue to be responsible for a metabolic function. Finally, we demonstrate the capacity of this approach to leverage metabolic functions of human cells and tissues using transcriptomic data from the Human Protein Atlas^11^ and show how the identification of metabolic tasks can be used to understand the organization of these biological entities into broader functional organ systems. Furthermore, using data from the Single-Cell Atlas of Adult Mouse Brain^12^, we show cell type specificity of several metabolic functions. Finally, we highlight the potential applications of this method to drive the discovery of new drug targets by identifying the main metabolic dysregulations associated with Alzheimer disease using single-cell transcriptomic data from the ROSMAP project^13^ (Religious Orders Study and Memory Aging Project).

## Results

### A framework to quantify a cell’s metabolic functions

Cells deploy diverse molecular functions to interface with their microenvironment, and adapt these as needed to cope with environmental changes. In metabolism, small modules of reactions can be defined as metabolic tasks (i.e., the generation of specific product metabolites given a defined set of substrate metabolites). The library of metabolic tasks a cell can sustain is encrypted in its genome and the capacity to modulate the activity of these tasks enable the cell’s adaptation to changing environment.

This concept of “metabolic tasks” has been previously used to evaluate the quality and capabilities of genome-scale metabolic models^9–11, 14–18^. However, these studies used various frameworks to define the cell’s capacity to sustain a metabolic task. Furthermore, the library of metabolic tasks used differed across studies in content and form. Thus, we first manually collated, curated and standardized existing metabolic task lists^9, 10^, resulting in a documented collection of 195 tasks covering 7 major metabolic activities of a cell (energy generation, nucleotide, carbohydrates, amino acid, lipid, vitamin & cofactor and glycan metabolism) (Figure 1, Supplementary Table 1). We further unified the formalism of the metabolic tasks and the associated computational framework for their use in the modelling context (details are presented in our earlier study^19^).

**Figure 1.**
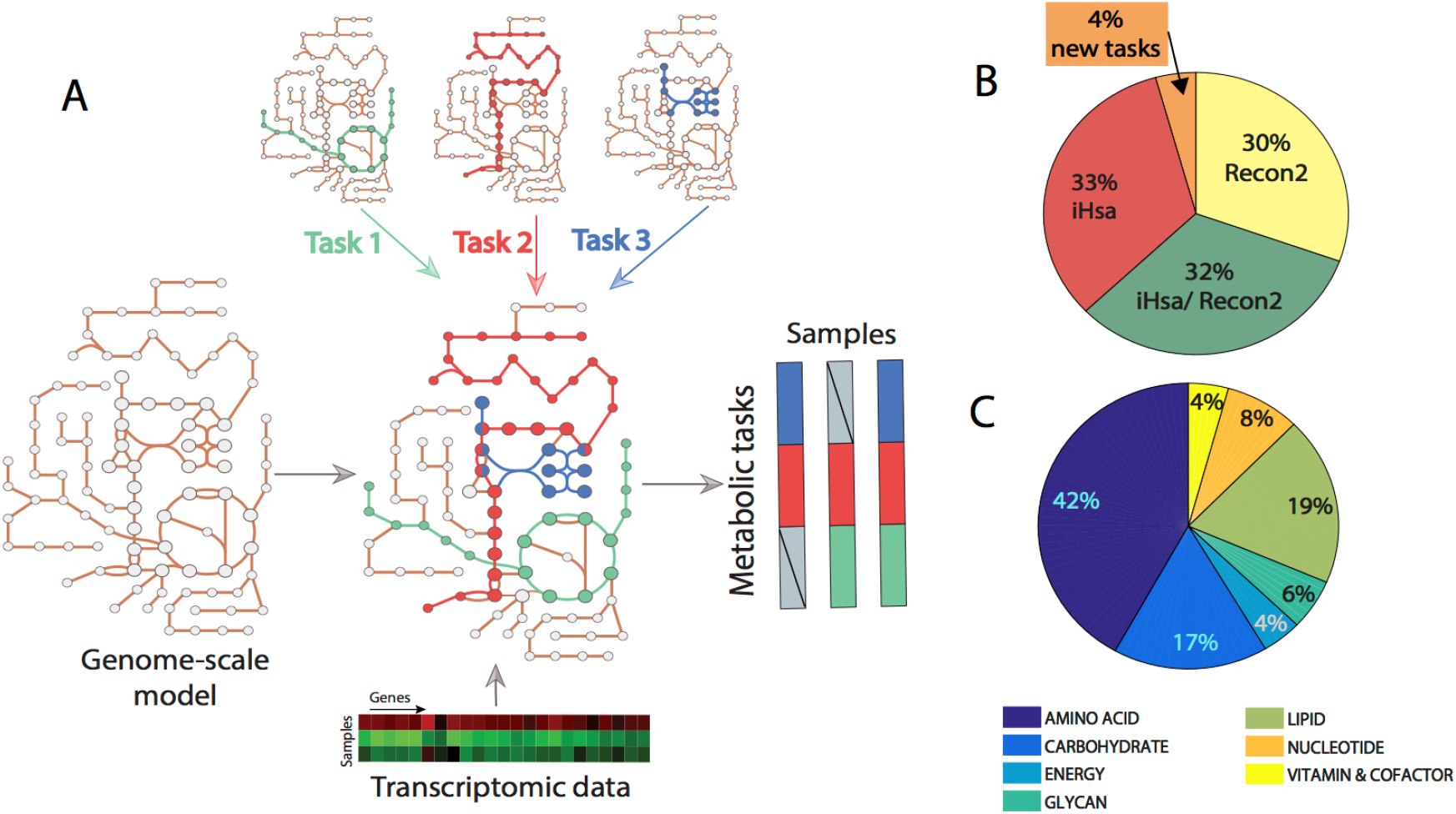
Genome scale metabolic models can be used to infer the activity of a defined list of metabolic functions. (A) Metabolic tasks are a modeling concept that we extend here to infer metabolic functions from transcriptomic data. (B) We curated and reconciled a collection of 195 tasks, derived in large part from earlier modeling studies (i.e., Recon 2 and iHsa). (C) The list of curated tasks covers seven main metabolic systems.

Here, we extend this concept beyond model benchmarking by developing a platform that quantifies a cell’s metabolic functions directly from transcriptomic data. To do this, genome-scale metabolic models are used to identify the list of reactions required to accomplish each metabolic task and, doing so, to identify the list of genes that may contribute to the acquisition of this metabolic function based on GPR rules (i.e., Gene Protein Reaction rules). The proposed computation of the metabolic score (i.e., relative activity of a metabolic task) relies first on the preprocessing of the available transcriptomic data and the attribution of a gene activity score for each gene^20^. We further select the genes responsible for the activation of each reaction required for a task using the GPR rules and average their activity to compute the metabolic task score (see Methods section for more details). Doing so, transcriptomic data can be directly used to quantify the relative activity of each metabolic function in a specific condition. Importantly, since gene lists are precomputed, no modeling background is required for the user.

### Metabolic tasks can leverage metabolic functions of human tissues

Each organ, tissue, and cell type in the human body carries out a distinct set of specific functions. The functions of each cell type are integrated to achieve the functions of each tissue, organ and organ system. Since there is no central database comprehensively describing the unique metabolic functions of different tissues, we used transcriptomic data from the Human Protein Atlas^11^ to quantify the metabolic functions of 32 tissues by using Recon 2.2^21^ as reference genome-scale model (Figure 2A, Supplementary Tables 2-3). We observed that >40% of the tasks are shared by all tissues (i.e., 79 tasks, Figure 2B), and within organ systems, even more tasks were shared (Figure 2C, Supplementary Table 4). To assess the significance to this common set of tasks, we collected a list of known housekeeping genes^22–25^. This list included 411 metabolic genes from Recon 2.2 (24.5% of all metabolic genes in Recon 2.2). Interestingly, we found that 97.5% of tasks shared by all the tissues are associated with at least one housekeeping gene. This included 277 housekeeping genes covered by metabolic tasks, which represent 67.4 % of all the Recon 2.2 housekeeping genes.

**Figure 2.**
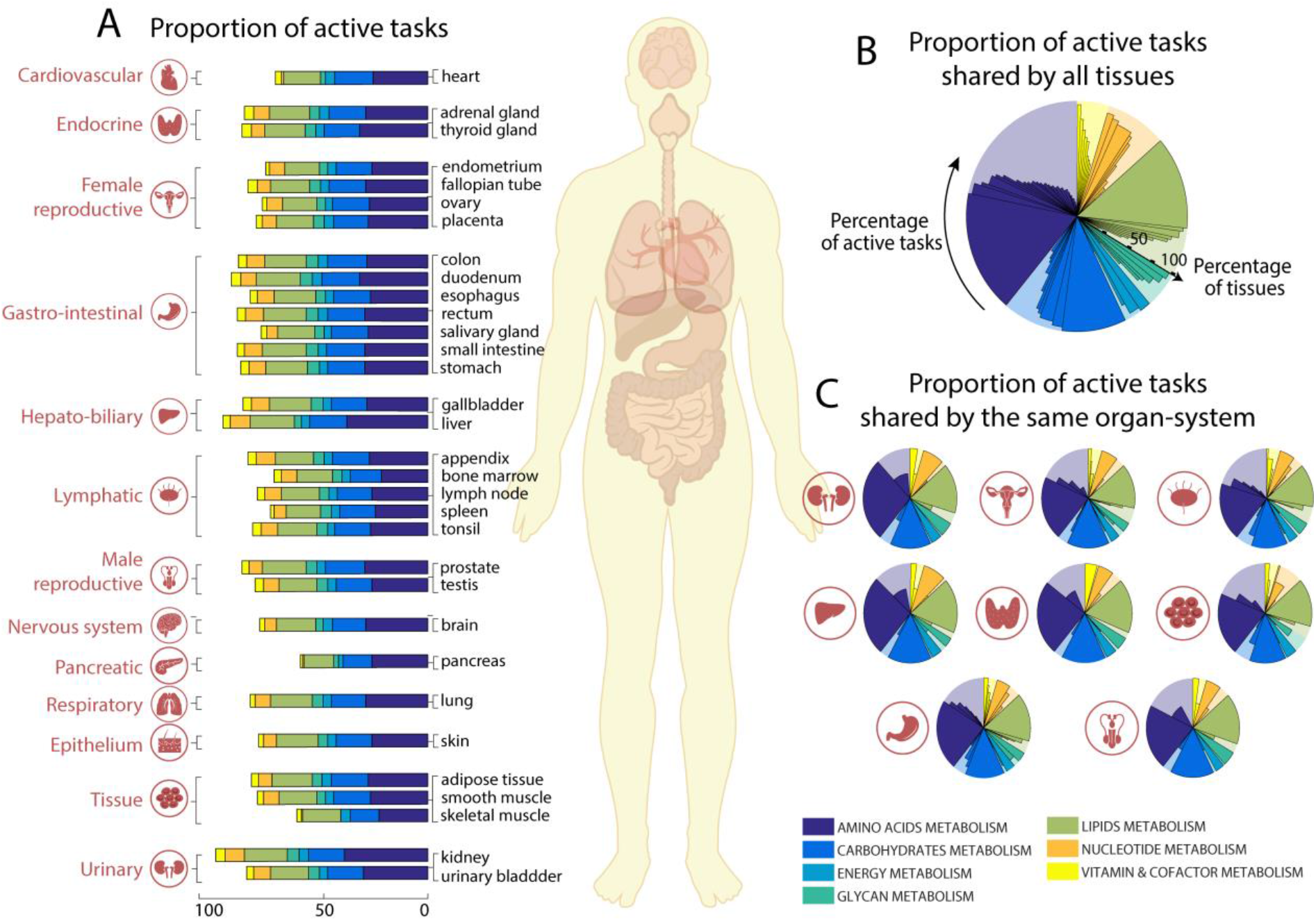
Metabolic tasks capture functional similarities between human tissues. (A) The proportion of tasks identified as active in the 7 major metabolic activities for each of the 32 tissues present in the Human Protein Atlas (Uhlen et al., 2015). (B) The percentage of active tasks that are shared by all tissues and (C) those shared within the same organ-systems (Supplementary Table 4). The background shaded color distribution represents the assignment of the 195 curated tasks to seven main metabolic systems.

### Metabolic tasks better capture within-tissue similarities than enrichment analysis

We further analyzed the similarities of metabolic tasks of tissues within the same organ systems. Specifically, we compared the similarities of tissues belonging to three different organ systems (i.e., female reproductive, gastrointestinal tract and lymphatic system) using either pathway enrichment analysis or the metabolic tasks (Supplementary Table 5, see Methods for more details). We found that the metabolic task approach significantly improves the grouping of tissues by organ system (Figure 3 A-B, Supplementary Figure 1).

**Figure 3.**
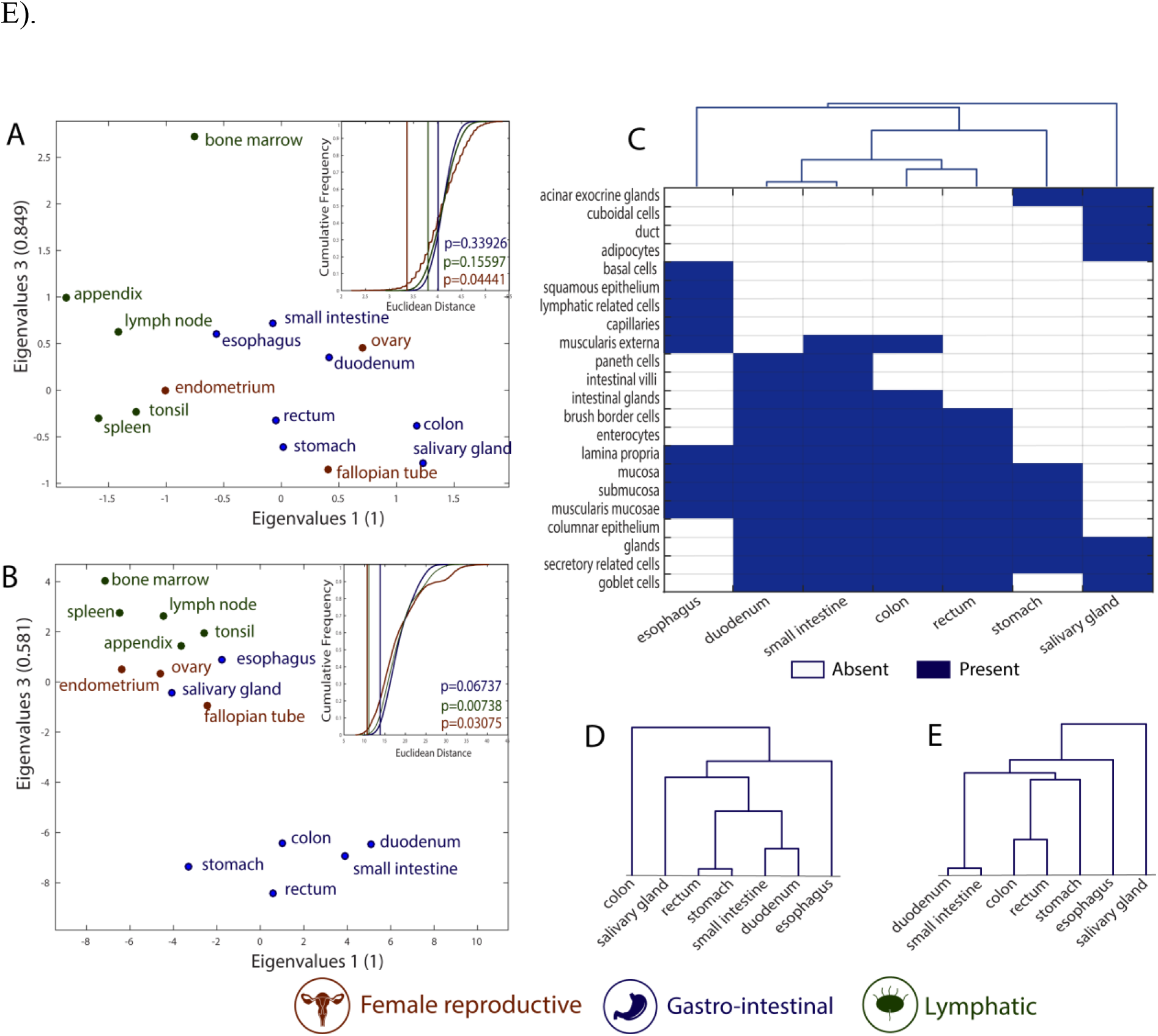
Metabolic tasks better capture the histological similarities of tissues. (A&B) Visual representation of the similarity between tissues computed based on (A) enrichment analysis and (B) the metabolic task approach using a principal coordinates analysis. Metabolic tasks cluster tissues into organ systems better than enriched pathways, as compared to the mean Euclidean distance for 100000 randomly selected groups with the same number of tissues (figure insets). The vertical lines are the mean Euclidean distance between tissues belonging to the same organ system and their empirical p-value (see Methods for more details). (C) Heatmap and hierarchical clustering of histological similarities between tissues of the gastrointestinal group. (D&E) Hierarchical clustering of similarities between tissues of the gastrointestinal group computed based on functional pathway enrichment (D) and the metabolic task approach (E)

The gastrointestinal system presents the lowest grouping significance for both approaches. Moreover, two tissues (i.e., esophagus and salivary gland) seem to be group outliers when the tissue similarity is assessed using the metabolic approach. Interestingly, these two tissues are histologically substantially different from the rest of the gastrointestinal system. Specifically, they are the only tissues without columnar epithelium. The salivary gland is the only tissue in this group having cuboidal cells in its epithelium, while the esophagus contains squamous epithelium (Figure 3C). The metabolic task approach better captures the histological distance between tissues belonging to the gastrointestinal system than enrichment analysis (Figure 3D-E).

### Metabolic task analysis captures tissue and cell specific functions

Some metabolic functions only occur in specific organs, tissues or cells. For example, taurine is the major constituent of bile secreted by the liver, and its biosynthesis also occurs in the kidney and brain^26^. Furthermore, taurine has been shown to play an important role in maintaining normal reproductive functions of mammals^27, 28^. Metabolic task analysis shows taurine synthesis in those known tissues and reproductive tissues (Figure 4A). Similarly, metabolic task analysis predicts that starch degradation occurs in the digestive tissues, consistent with the reported localization^29^. Thus, the analysis can capture tissue-specific metabolism.

**Figure 4.**
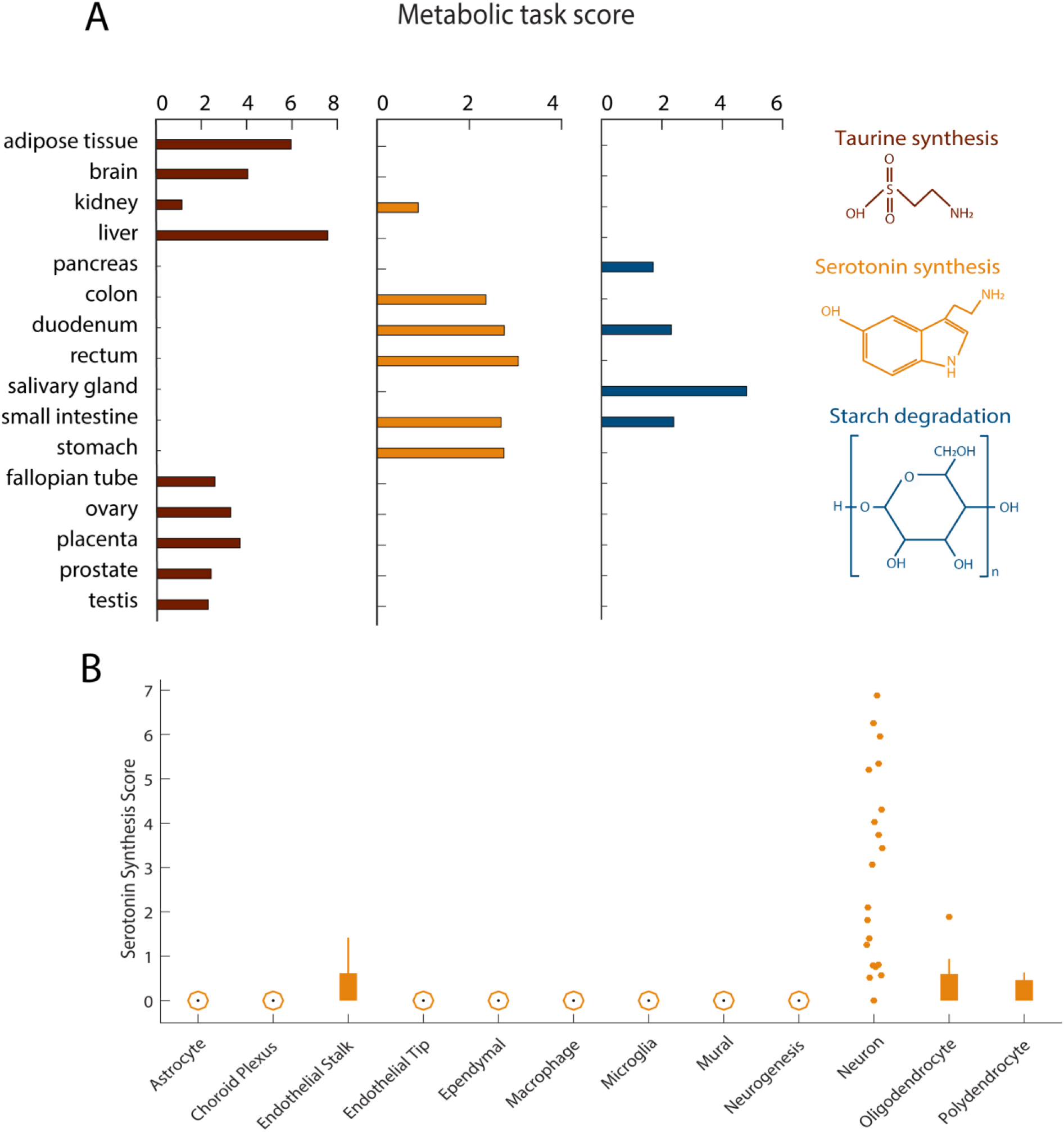
Metabolic specificities of tissues and brain cells. (A) Metabolic task scores associated with the synthesis of taurine and serotonin and the degradation of starch. Note, the figure only presents the 16 tissues where these tasks have been predicted. (B) Score associated with the synthesis of serotonin for 12 different brain cell types.

Serotonin biosynthesis is similarly accurately predicted to be synthesized in the gastro-intestinal tract. However, the method does not predict its known synthesis by the brain^30^. This can be expected as serotonergic neurons are localized to the raphe nuclei, whereas the bulk brain transcriptomic data in the HPA RNA-Seq were sampled from cerebral cortex^11^. Thus, we used the metabolic task approach on single-cell RNA-Seq data of the adult mouse brain^12^ (Supplementary Tables 6- 7), and found that serotonergic neurons can be successfully identified (Figure 4B).

### Metabolic task analysis captures the differences between brain cell types

The human brain is a metabolically demanding organ consisting of diverse cell types, each one with unique metabolic capabilities. While some metabolic interchanges between brain cell types are well-known (e.g., glutamate-glutamine shuttle between neurons and astrocytes), it remains many open questions concerning the specific contribution of each cell-type in brain function. In this context, we used single cell RNA-Seq data from adult mouse brain^12^ to assess the main metabolic features that differentiate astrocytes, neurons and oligodendrocytes (Figure 5A, see Methods for more details). The metabolic task approach clearly differentiates the three cell-types and details their metabolic specialization (Figure 5B-C). Our analysis confirms previously known specific metabolic features such the evidence that astrocytes fuel the glutamate-glutamine shuttle^31^ (Figure 5B) and that oligodendrocytes are likely the primary source of creatine in the brain^32^ (Figure 5C). Interestingly, there has been a debate as to if oligodendrocytes serve as sources of glutamine synthesis^33^ in the glutamate-glutamine shuttle. Our analysis of single cell RNA-Seq clearly supports the hypothesis (Figure 5B, Supplementary Figure 3D).

**Figure 5.**
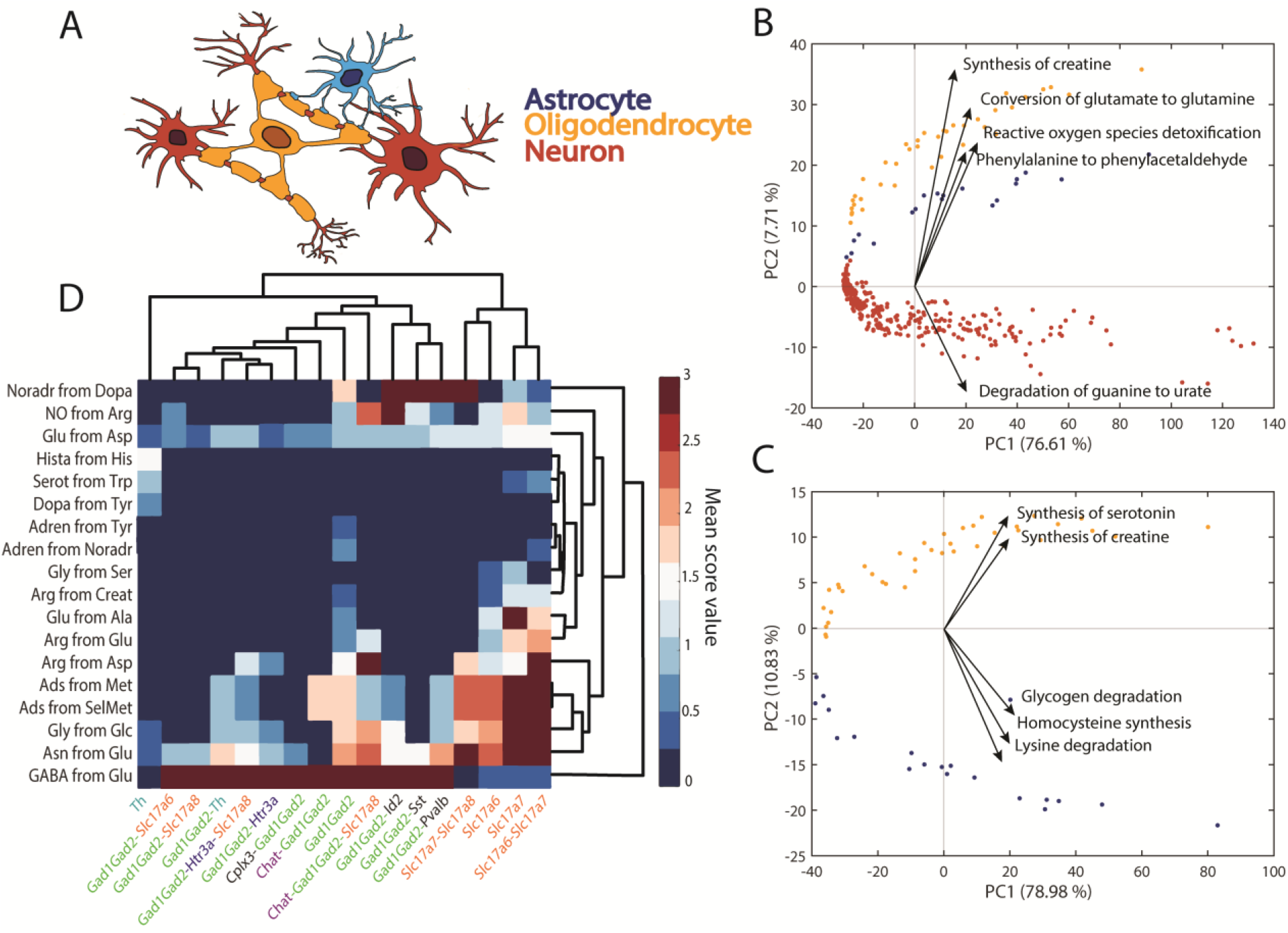
Metabolic differences between astrocytes, neurons and oligodendrocytes. (A) Schematic representation of spatial connection between astrocytes, neurons and oligodendrocytes. (B) PCA component scores for the 3 different cell types (astrocytes, neurons, oligodendrocytes) and the 5 dominant tasks in the second principal component. (C) PCA component scores for only 2 cell types (astrocytes, oligodendrocytes) and the 5 dominant tasks in the second principal component. (D) Heatmap of metabolic tasks score mean values associated with the synthesis of main neurotransmitters in the context of the gene markers for different neuron types.

To analyze the capacity of this method to be used to resolve open questions, we also created a new set of tasks specific to neurotransmitters synthesis (Supplementary Table 8). We compared the expression of these tasks with respect to the type of gene markers used to differentiate the single cells. We can observe that each set of gene markers used to identify the different clusters of neurons in the single-cell atlas of adult mouse brain^12^ are associated with specific neurotransmitter patterns. Specifically, the Slc17 gene family is associated with the non-expression of GABA neurotransmitter presumably corresponding to glutamatergic neurons. Contrarily, all the neurons identified using Gad family genes markers are associated with a high GABA synthesis presumably corresponding to GABAergic neurons^12^. Interestingly, tyrosine hydroxylase (Th) is often used as a marker of dopaminergic neurons^34^. We can observe that the neurons identified with this gene are the only ones presenting the synthesis of dopamine.

### Metabolic task analysis highlights the main metabolic dysregulations in Alzheimer’s disease

Alzheimer’s disease is a neurodegenerative disorder affecting millions of people, but to date we lack a true cure. Despite decades of research into the disease, many questions remain regarding the molecular basis of its progression. However, increasing evidence suggests metabolic dysfunction may contribute to nervous system degeneration^35–37^. Whether metabolic alterations are the cause or the consequence of the pathogenesis remains unclear, but it stands to reason that metabolic pathways may themselves contain potential targets for future therapies^38^. In this context, we used single-cell RNA-Seq data from the ROSMAP project^13^ to elucidate the main metabolic dysregulations associated with Alzheimer’s disease. To this end, we clustered the excitatory neuron samples and identified the tasks that were active in more than 50% of the dataset. Only three metabolic tasks correspond to this criterion: the conversion of phosphatidyl-1D-myo-inositol to 1D-myo-inositol 1-phosphate, the synthesis of tetrahydrofolate and the synthesis of “Tn_antigen” (i.e., Glycoprotein N-acetyl-D-galactosamine). We further used them to divide the samples into 8 metabolic clusters depending on the combination of their activity in each sample (Figure 6A-B, see Methods for more details).

**Figure 6.**
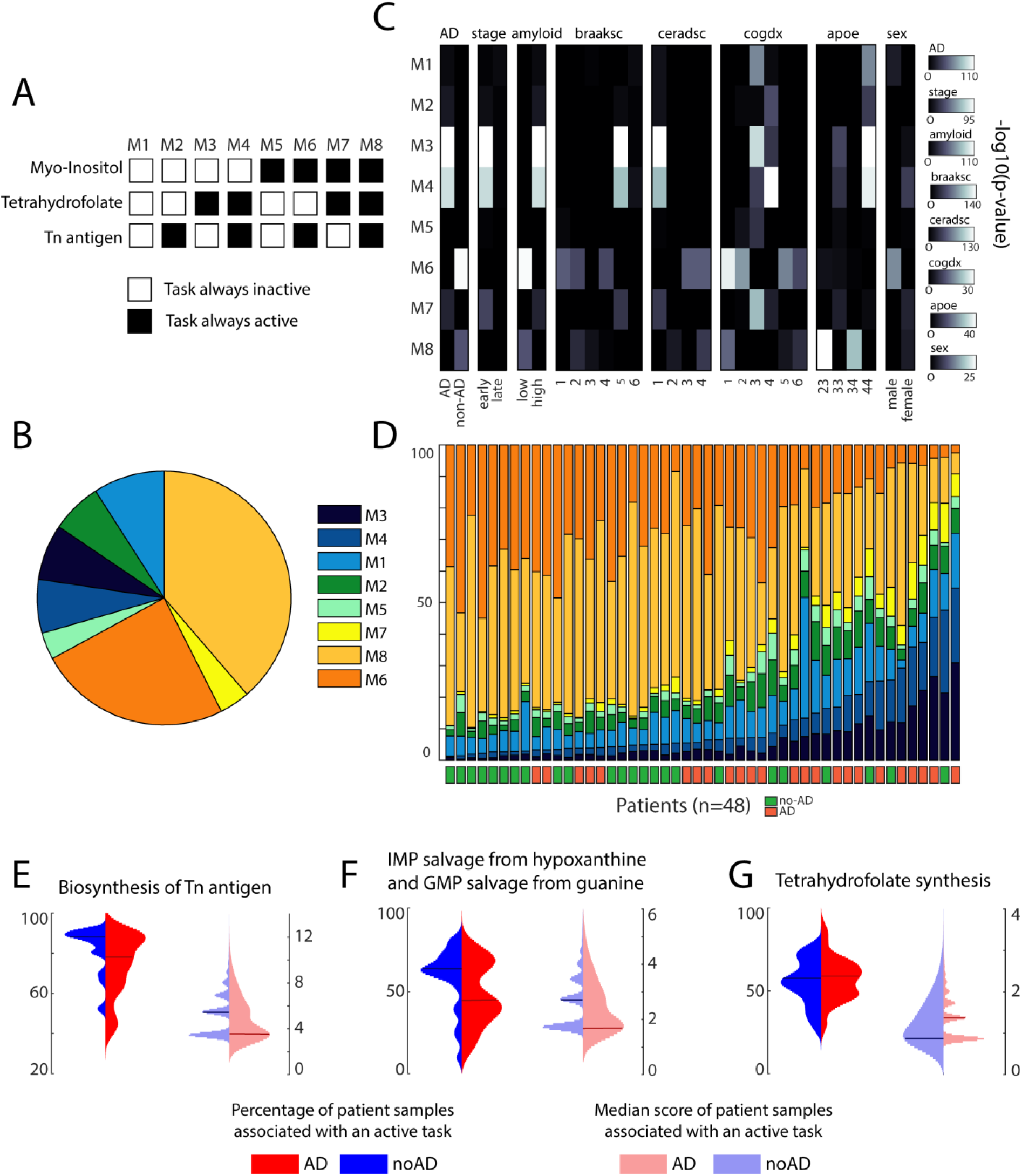
Metabolic subpopulations of excitatory neurons and their link with Alzheimer’s disease. (A) The single-cell transcriptomic dataset was clustered into 8 metabolic subpopulations with distinct patterns of activity for 3 metabolic tasks. (B) Percentage of the representation of each metabolic cluster within the dataset. (C) Enrichment analysis (one-tailed Fisher’s exact test) within each metabolic cluster of clinic-pathological variables^39^. (D) Percentage of samples of each metabolic cluster from each patient and their associated Alzheimer’s diagnosis. (E-G) Expression patterns of the metabolic tasks (left - percentage of patient samples associated with an active task and right - related median score) presenting a dysregulated activity across groups of patients with different diagnosis for Alzheimer’s disease (blue – patient without Alzheimer and red – patients with Alzheimer). The horizontal lines represent the median of the distribution.

For each metabolic cluster, we tested their associations with pathological traits using a one-tailed Fisher test (Figure 6C) and observed that specific metabolic clusters were enriched in samples associated with either Alzheimer’s pathology (clusters M3 and M4) or no pathology (cluster M6). Interestingly, we were able to group the 48 patients from the dataset depending on their disease prognostic with 75% accuracy by sorting them with respect to the proportion of their samples in M3 and M4 (Figure 6D). Note that we applied the clustering approach and subsequent trait enrichment analysis to the 6 major cell types identified in the original study presenting this dataset^39^ and we did not find such strong correlation for the other brain cell types (Supplementary Table 9).

To better understand, the metabolic functions differentiating the 8 clusters, we computed the median of the combined metabolic task score (i.e., score in its binary version multiplied with the continuous one) and observed that only 13 tasks presented a median score different than zero in a metabolic cluster. We further used these identified tasks to investigate their expression patterns (i.e., percentage of patient samples associated with an active task and related median score) across the groups of patients presenting or not a positive diagnosis for Alzheimer’s disease. Very distinctive median score distribution depending on the Alzheimer’s diagnosis are observed for 4 tasks previously highlighted in the literature as being implicated in the Alzheimer’s disease (Figure 6E-G): the synthesis of Tn antigen^40, 41^ (glycoprotein N-acetyl-galactosamine), the synthesis of tetrahydrofolate^42^ and the salvage of IMP and GMP^43^. While the other metabolic tasks identified do not present distinctive patterns at the level of the median score distribution, we can observe that healthy subjects often present a higher percentage of samples for which these tasks are active (Supplementary Figure 5). The results imply that an overall deficiency of these metabolic activities is observed in Alzheimer’s disease patients. Interestingly, some of these dysregulated metabolic tasks have been observed in previous studies, such as the pyridoxal phosphate synthesis^44^, the presence of the thioredoxin synthesis^45^, the fructose degradation^46^ and the conversion of myo-inositol^47^, while the others have not been specifically investigated. In this context, the metabolic dysregulations identified with our approach might be of interest to drive the discovery of new potential drug targets.

## Discussion

Here, we present an approach to predict the activity of hundreds of metabolic functions from transcriptomic data. This framework enables the comprehensive quantification of the propensity of a cell line or tissue to express a metabolic function, thereby facilitating phenotype-relevant interpretation of these complex data types. We used multiple omics datasets to highlight the power of our approach to quantify metabolic functions from organ systems to single cells and demonstrated its advantages compared to functional enrichment analysis approaches that are widely used to connect measured gene expression changes to phenotypes^1, 2^.

Enrichment analyses provide invaluable insights but they have a limited capacity to describe how the changes in gene expression impact pathway functions. Indeed, enrichment approaches only take in gene categories based on known associations but usually do not leverage the biological mechanisms. This can be exemplified in that a group of genes might be in the same pathway gene set without contributing, to the same extent, to the metabolic pathway’s specific function. Thus, while enrichment analyses provide the molecular context to facilitate interpretation, it remains challenging to mechanistically link changes in the enriched pathways to the regulation of the biological processes and their implications in a specific disease.

Our framework helps address these limitations by analyzing gene expression with a network-based approach and therefore accounts for gene dependencies in pathway functions. This approach integrates omics datasets into pathways from computational models to quantitatively describe the genotype-phenotype relationship. The analysis of gene expression data with genome-scale systems biology models is well established and can provide deep mechanistic insights into the metabolic capabilities of a cell and/or a tissue. Indeed, Uhlen et al.^11^ used a network-based approach and the concept of metabolic tasks to construct tissue-specific metabolic networks. The approach enforced the activity of tissue-specific metabolic tasks into each model to capture cellular functionalities known to occur in all cell types. Doing so, they also found metabolic housekeeping functions shared across all tissues and showed similarities between metabolic activities across tissues in the same organ systems. Unfortunately, the construction and analysis of such computational models is a complex and difficult task requiring expert knowledge of the tissues and modeling framework^19, 20, 48^. To overcome this problem, our framework successfully combines the capacity to provide mechanistic insights of network based approaches and the simplicity of enrichment analyses. To further facilitate adoption of the approach, we created a web-based CellFie module that has been integrated into the list of tools available in GenePattern^49^ (www.genepattern.org, see Methods for more details).

Our tasks cover hundreds of mammalian metabolic functions, but can be easily extended to diverse organisms and more cellular functions captured in systems biology models of metabolism, transcription, translation, signaling, etc. For example, genome-scale metabolic networks exist for hundreds of organisms. A community standard for metabolic tasks will facilitate efforts to build an extensive resource of metabolic and cellular functions. Furthermore, while the inclusion of other biological processes (e.g., transcription, translation) may require different types of models^50, 51^, our approach only requires gene information and therefore can easily be formulated into our framework. Finally, future work will investigate contributions from different isoenzymes within each metabolic task, since different cells and tissues can present the same metabolic reactions, but using different isoenzymes with different activities^11^.

This variation in enzyme usage may underlie adaptations of metabolism to biological perturbation such as a disease.

In conclusion, this framework provides a new analysis approach combining the best of existing methodologies (i.e., network- and knowledge-based functional analysis). This might, one day, enable the complete deconstruction of the molecular basis of any biological system based on a simple omics data analysis.

## Methods

### Gene expression data and models

RNA-Seq data for the 32 human tissues were downloaded from the Human Protein Atlas^11^. The brain single-cell transcriptomic datasets come from Cell Atlas of Adult Mouse Brain^12^ and the ROSMAP project^13^ (Religious Orders Study and Memory Aging Project). The ROSMAP data can be requested at www.radc.rush.edu. The models used to assess the metabolic task is iMM1415^52^ for the single cells from adult mouse brain and Recon2.2^21^ for human tissues and the human brain cell types of Alzheimer patients.

### Preprocessing of gene expression data

We processed the gene expression data to attribute a gene activity score for each gene and define which genes are active in each cell or tissue. A gene is defined as active in a sample if its expression value is above a threshold defined for this gene within the dataset considered. The threshold of a gene is defined by the mean value of its expression over all the samples coming from the same dataset with exceptions that the threshold need to be higher or equal the 25^th^ percentile of the overall gene expression value distribution and lower or equal to the 75^th^ percentile. The gene score is computed as follow:

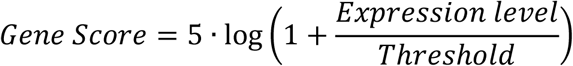

These gene scores are mapped to the models by parsing the GPR rules associated with each reaction. The gene score for each reaction is selected by taking the *minimum* expression value amongst all the genes associated to an enzyme complex (AND rule) and the *maximum* expression value amongst all the genes associated to an isozyme (OR rule)^53^. Note that we have recently benchmarked the influence of preprocessing methods on the definition of the set of active genes and observed that this parameter combination presented the best performance^20^.

### Curation of metabolic tasks

The curation was done by first taking the union of previously published lists of metabolic tasks^9, 10^. We removed duplicated tasks and lumped tasks that rely on the description of similar metabolic functions. Each remaining task without strong biological evidence was removed. We also created 9 new tasks that were essential for the acquisition of already described metabolic functions (i.e., intermediate biosynthetic steps for the acquisition of other tasks). Doing so, we obtained a collection of 195 tasks associated with 7 systems (energy, nucleotide, carbohydrates, amino acid, lipid, vitamin & cofactor and glycan metabolism). For each task, we provided its original source (Recon and/or iHsa) and comments on the biological evidence of this metabolic function (Supplementary Table 1).

### Inference of metabolic tasks from transcriptomic data

We developed a computational framework for attributing a score to each metabolic task in order to extend the application of the concept beyond the model benchmarking scope. If a task successfully passes in a model, one can compute the list of reactions associated with the task and, doing so, access to the list of genes that may contribute to the acquisition of this metabolic function based on the GPR rules. To this end, we used the parsimonious Flux Balance Analysis (pFBA) algorithm to define the set of reactions and associated genes required to pass a task within a specified model^54^. Thanks to the availability of this information, metabolic functions can now be directly assessed from transcriptomic data. The proposed computation of a metabolic score relies first on the preprocessing of the available transcriptomic data and the attribution of a gene activity score for each gene (see associated Methods section). We further used the GPR rules associated with each reaction required for a task to decide which gene will be the main determinant of the enzyme abundance associated with this reaction and attribute the corresponding gene activity level. Therefore, each reaction involved in a task is associated with a reaction activity level (RAL) that corresponds to the preprocessed gene expression value of the gene selected as the main determinant for this reaction. We also computed the significance of each gene selected with regard to its overall use in the observed condition. Actually, some genes will be mapped to multiple reactions (e.g. promiscuous enzyme). Therefore, we assume that there may exist some competition between the reactions using this gene. We define the significance of a gene (S) by its specificity for a reaction by the inverse of the number reactions in which this gene is used as the main determinant. Finally, the metabolic score can be computed as the mean of the product of the activity level of each reaction with the significance of its associated gene:

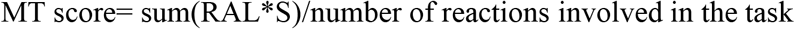

MT score provides a relative quantification of the activity of a metabolic task in a specific condition based on the availability of data for multiple conditions. Indeed, it has been shown that some important housekeeping genes always present very low expression value. Therefore, a metabolic function that will completely rely on this set of genes will always result in a low MT score. Contrarily, some tasks can be associated with gene presenting very high expression levels. Therefore, MT scores cannot be compared across tasks but only across samples. To partly overcome this problem, we also propose this scoring approach in its binary version to determine whether a metabolic task is active or not based on a gene expression profile. To this end, the MT score no longer takes into account the significance of gene determinant for each reaction but is just computed as the mean of the reaction activity levels. Doing so, a metabolic task will be considered as active if its MT score in its binary version has a value superior to 5log(2).

### Assessment of tissue similarities

We computed the scores of the 195 metabolic tasks in their continuous version based on the transcriptomic data available for 32 different tissues in the Human Protein Atlas^11^ dataset using Recon2.2 as reference genome-scale metabolic model^21^. These scores were used to compute the Euclidean distance between each tissue. We associated each tissue to an organ system (Supplementary Table 4) and computed the average Euclidean distance between tissues belonging to the same organ system. Note that, we only considered organ systems presenting more than two tissues within the same group (i.e. Female Reproductive, Lymphatic and Gastrointestinal). To compute the significance of our results, we generated the mean Euclidean distance for 10000 randomly selected groups with the same number of tissues and computed the exact p value (i.e. proportion of random distance lower than the observed distance) associated to each organ system. We performed the same type of analysis based on the results of functional pathways enrichment obtained using Enrichr^55^ (Supplementary Table 5) and presented a comparison in Figure 4. Note that we also performed this analysis using the metabolic scores when computed in their binary version (Supplementary Table 3 and Supplementary Figure 1). The histological information used in the assessment of tissue similarities has been collected from the microscopy images and associated description available in the Human Protein Atlas^11^.

### Principal component analysis for differentiating brain cell-types

A matrix representing the metabolic function scores for 3 brain cell types (i.e., astrocytes, neurons and oligodendrocytes) was constructed by multiplying the metabolic task scores computed in their continuous version (Supplementary Table 6) with the ones in their binary version (Supplementary Table 7). A PCA analysis on this matrix was conducted. As this analysis did not enable the differentiation between astrocytes and oligodendrocytes, we performed a subsequent similar PCA analysis by only using the samples related to these specific cell-types.

### Clustering of excitatory neurons samples from the ROSMAP project

We clustered the samples identified as excitatory neurons by identifying the tasks that were actives in more than 50% of the dataset. This threshold has been set with respect to the percentage of excitatory neurons samples associated with a positive diagnosis of Alzheimer’s disease (i.e., 51,2%). Only three metabolic tasks correspond to this criterion: the conversion of phosphatidyl-1D-myo-inositol to 1D-myo-inositol 1-phosphate, the synthesis of tetrahydrofolate synthesis and the synthesis of Tn_antigen (Glycoprotein N-acetyl-D-galactosamine. We further used them to divide the samples into 8 metabolic clusters depending on the combination of their activity in each sample (Figure 6A-B). Note that prior to this choice, other clustering methods have been investigated. Our first approach was using a k-means clustering method. To this end, we used the percentage of coordinates that differ (hamming distance) in the binary matrix of the metabolic task score (active vs non-active) and the matlab function k-means with 10 replicates. To identify the appropriate number of clusters to separate the data, we computed the within-cluster sum of square distance (wws) and the average silhouette value by iteratively increasing the number of clusters from 1 to 15. This approach also led to the identification of 8 metabolic clusters that were displaying the same metabolic dysregulations. In order to ensure the reproducibility of the results presented, we preferred to use a more straightforward clustering method.

We compared the metabolic clusters obtained with our approach to the clusters identified in a publication^39^ using the ROSMAP data (Supplementary Figure 4). We can observe that the metabolic clusters M3 and M4 are only enriched in clusters Ex2 and Ex4 who were identified as highly correlated with Alzheimer’s pathological traits in the reference publication. The same observation can be done with M6 metabolic cluster and Ex6, the cell type cluster identified as highly correlated with patients without Alzheimer’s disease.

### Web-based CellFie module to perform analysis

We created a web-based CellFie module that has been integrated into the list of tools available in GenePattern^49^ (www.genepattern.org). A tutorial explaining how to run CellFie as a GenePattern module is available on the wiki section of the github repository: https://github.com/LewisLabUCSD/CellFie. This repository includes the source code of the computation framework that are running on Matlab and require the installation of the Cobra Toolbox^56^. It also includes a tutorial to visualize the output results of CellFie on metabolic maps using Escher^57^.

## Author contribution

AR, and NEL designed the study, conducted the analyses, and wrote the paper. AWTC, AR, JMG, CJ, JKL and SL curated the list of metabolic tasks. AR, ATW, BPK, DB, EFJ, JPM, KR, TB and TR developed the CellFie webtool. LH and CT organized the code repository in Github and integrated it with Artenolis. HM, JL and ZL created the tutorial to visualize the results obtained with the CellFie webtool on Escher maps. All authors have read and approved the work.

## Acknowledgements

This work was supported by generous funding from NIGMS to NEL (grant no. R35 GM119850), a LIFA fellowship to AR, grants for the support of GenePattern to JPM (R01 GM074024 and U24 CA194107), grants for ATW (NLM T15LM011271, Training Fellowship from UC San Diego Cancer Cell Map Initiative - NCI U54 CA209891). The Reproducible Research Results (R3) team of the Luxembourg Centre for Systems Biomedicine is acknowledged for supporting the project and promoting reproducible research. Single-cell transcriptomic data from the ROSMAP project were provided by the Rush Alzheimer’s Disease Center, Rush University Medical Center, Chicago. Data Collection was supported through funding by NIA grants P30AG10161, R01AG15819, R01AG17917, R01AG30146, R01AG36836, U01AG32984, U01AG46152, U01AG61356, the Illinois Department of Public Health, and the Translational Genomics Research Institute. Images of human tissues used in Figure 2 and images of brain cell types used in Figure 5A are adapted from work created by Freepik.com.

