## Supplementary Figures for "What does your cell really do? Model-based assessment of mammalian cells metabolic functionalities using omics data"

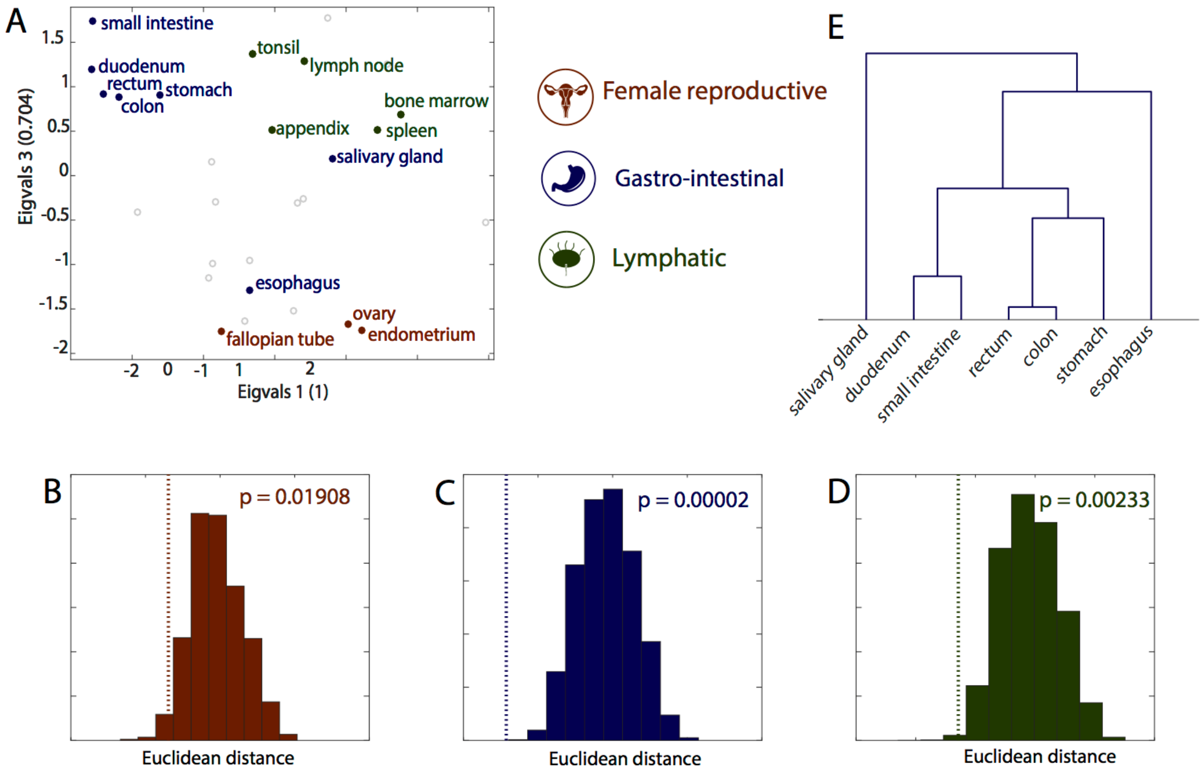


**Supplementary Figure 1** – Metabolic task in its binary version still better captures tissue groups at organ system level. (A) Visual representation of the similarity between tissues using a principal coordinates analysis (B, C & D). Significance of tissue grouping at the organ system level. Distribution of mean Euclidean distance for 100000 randomly selected group with the same number of tissues. The dotted line is the mean Euclidean distance between tissues belonging to the same organ system and it associated pvalue (see Methods for more details). (E) Hierarchical clustering of similarities between tissues of the gastrointestinal group.

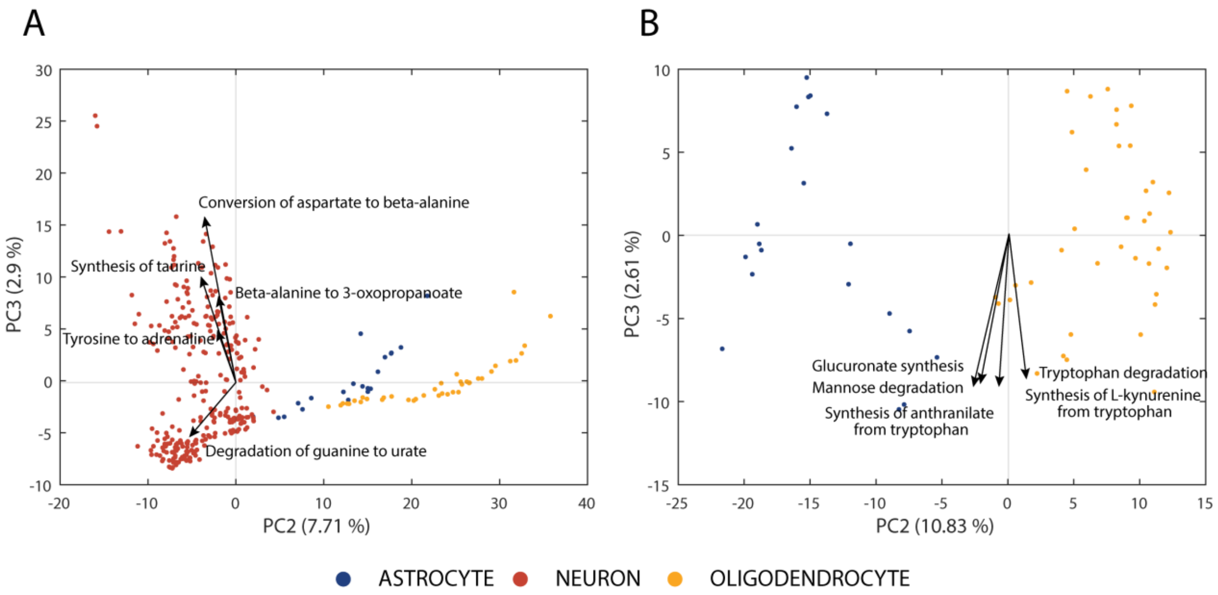


**Supplementary Figure 2. Principal Components Analysis -** (A) Component scores plot of the PCA performed using the 3 different cell types (astrocytes, neurons, oligodendrocytes) and the 5 tasks influencing the most the third principal component. (B) Component scores plot of the PCA performed using only 2 cell types (astrocytes, oligodendrocytes) and the 5 tasks influencing the most the third principal component.

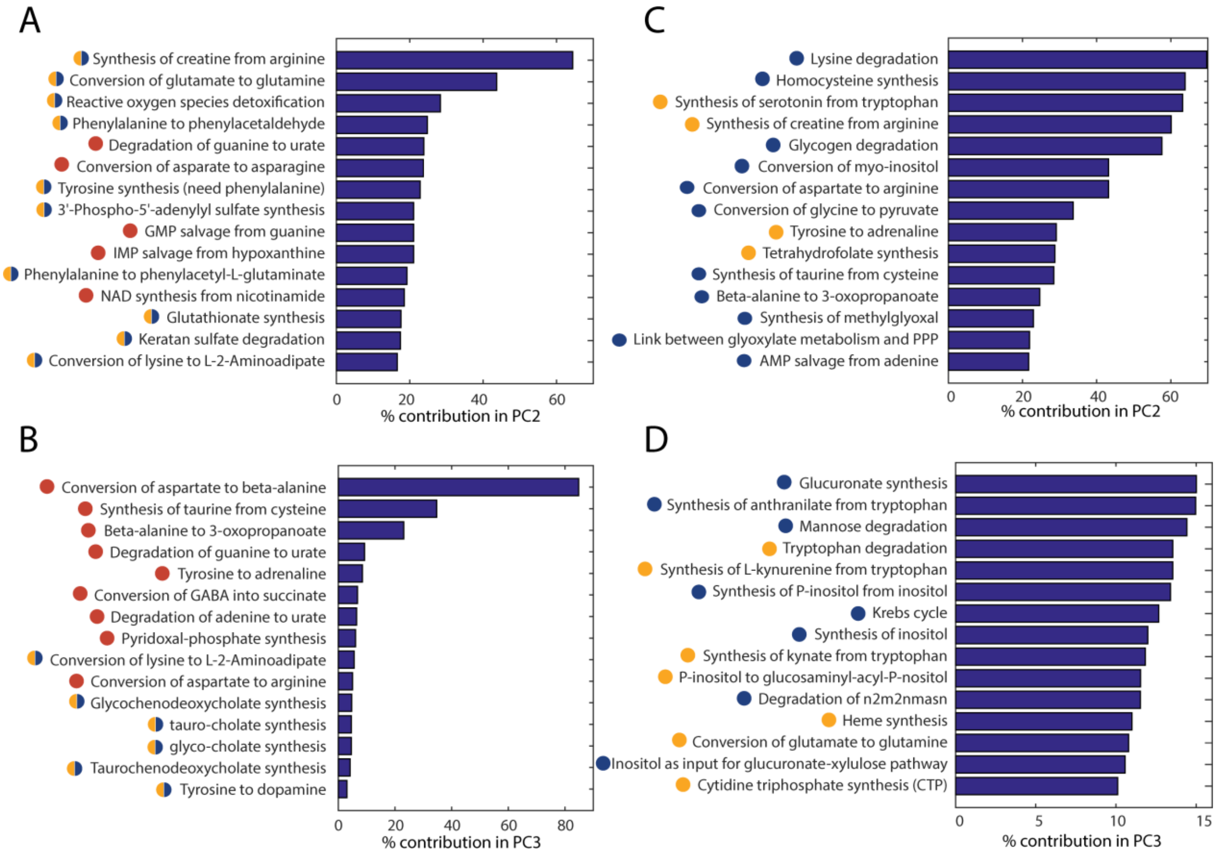


**Supplementary Figure 3. PCA loadings -** Contribution of the task in the principal components 2 (A) and 3 (B) in the analysis performed using the three different cell types (astrocytes, neurons, oligodendrocytes) and in the principal components 2 (C) and 3 (D) in the analysis performed using the samples associated to astrocytes and oligodendrocytes. Note that we did not plot the contribution of variables in PC1 has this component was not informative to differentiate the cell-types. The color bullets represent the direction of the loadings for the separation between the different cell-types (blue = astrocytes, yellow = oligodendrocytes and red = neurons).

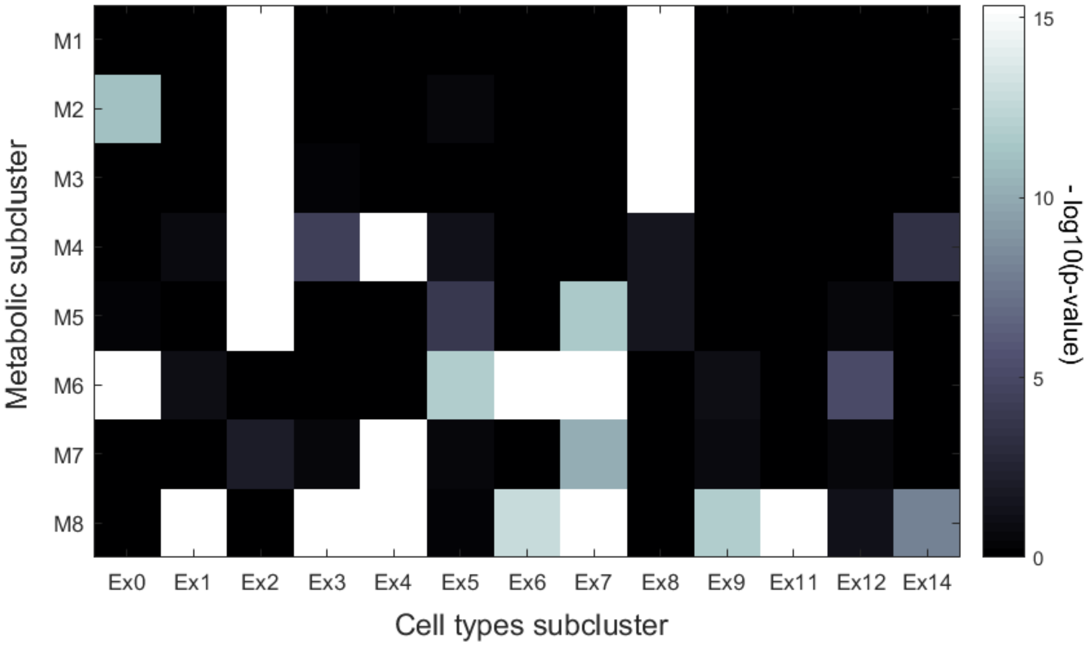


**Supplementary Figure 4.** Enrichment analysis (hypergeometric test) within the cell types subclusters identified in the original reference paper (columns) of the metabolic subclusters identified with our approach (rows).

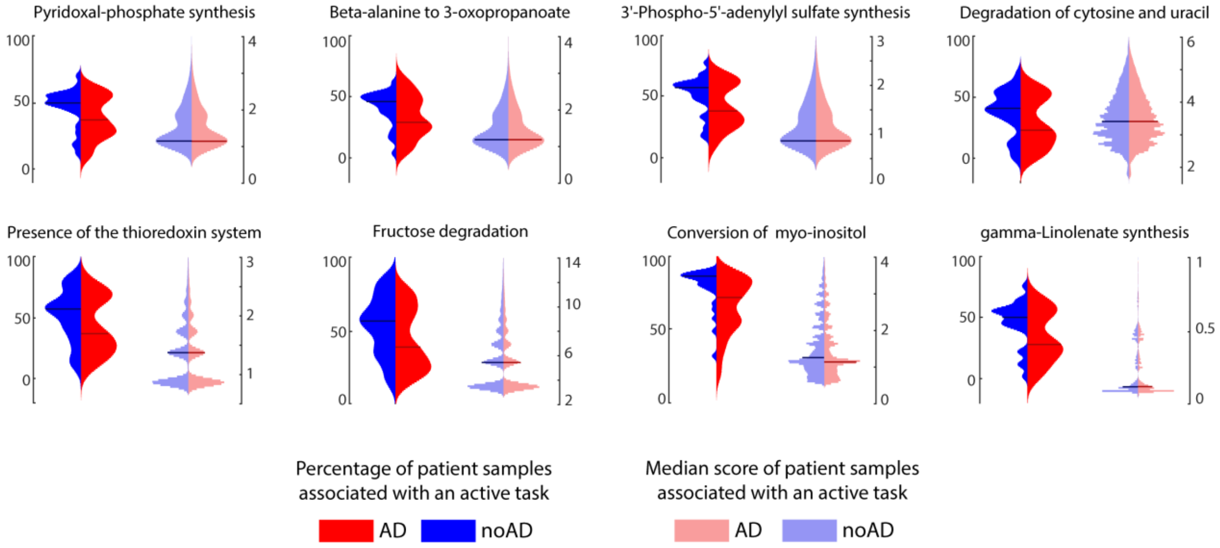


**Supplementary Figure 5.** Expression patterns of the metabolic tasks (left - percentage of patient samples associated with an active task and right - related median score) presenting a dysregulated activity across group of patients with different diagnosis for Alzheimer's disease (blue – patient without Alzheimer and red – patients with Alzheimer). The horizontal lines represent the median of the distribution.
